# Use of standard U-bottom and V-bottom well plates to generate neuroepithelial embryoid bodies

**DOI:** 10.1101/2021.12.17.473219

**Authors:** David Choy Buentello, Lena Sophie Koch, Grissel Trujillo-de Santiago, Mario Moisés Alvarez, Kerensa Broersen

## Abstract

The use of organoids has become increasingly popular recently due to their self-organizing abilities, which facilitate developmental and disease modeling. Various methods have been described to create embryoid bodies (EBs) generated from embryonic or pluripotent stem cells but with varying levels of differentiation success and producing organoids of variable size. Commercial ultra-low attachment (ULA) V-bottom well plates are frequently used to generate EBs. These plates are relatively expensive and not as widely available as standard concave well plates. Here, we describe a cost-effective and low labor-intensive method that creates homogeneous EBs at high yield in standard V- and U-bottom well plates by applying an anti-adherence solution to reduce surface attachment, followed by centrifugation to enhance cellular aggregation. We also explore the effect of different seeding densities, in the range of 1 to 11 ×10^3^ cells per well, for the fabrication of neuroepithelial EBs. Our results show that the use of V-bottom well plates briefly treated with anti-adherent solution (for 5 min at room temperature) consistently yields functional neural EBs in the range of seeding densities from 5 to 11×10^3^ cells per well. A brief post-seeding centrifugation step further enhances EB establishment. EBs fabricated using centrifugation exhibited lower variability in their final size than their non-centrifuged counterparts, and centrifugation also improved EB yield. The span of conditions for reliable EB production is narrower in U-bottom wells than in V-bottom wells (i.e., seeding densities between 7×10^3^ and 11×10^3^ and using a centrifugation step). We show that EBs generated by the protocols introduced here successfully developed into neural organoids and expressed the relevant markers associated with their lineages.

We anticipate that the cost-effective and easily implemented protocols presented here will greatly facilitate the generation of EBs, thereby further democratizing the worldwide ability to conduct organoid-based research.

## Introduction

Embryoid bodies (EBs) have garnered great interest in recent years as a powerful tissue engineering platform due to their self-reproducing and multipotent differentiation capabilities. EBs are precursors to organoids, whose self-organizing properties allow their exploitation as developmental and disease models [1–3]. While procedures exist for organoid differentiation, the success of the differentiation process relies heavily on the ability to control certain variables involved in the formation of EBs [4,5]. For example, having a spherical form is a prerequisite for most EB protocols, as this standardizes the gradient profiles that contribute to homogeneity [6]. However, arguably, the most important parameter when considering successful organoid differentiation is the initial diameter of the EB itself. Variation in diameter size severely affects differentiation, as increases in diameter result in longer diffusion times, thereby reducing the number of cells in the inner parts of the EB that have immediate contact with nutrients and differentiation factors [7,8]. Many traditional protocols cite EB diameter sizes ranging from 150 to 600 μm [9,10]; however, these sizes are generally not standardized. Consequently, current protocols result in EBs exhibiting wide variability in their diameters (i.e., 50 μm and higher). Having better control of the initial cell aggregate diameter would improve the differentiation of EBs into functional organoids. Previous studies have shown differentiation yields greater than 90% with improved culture methods [11,12]. A fabrication process that enhances EB homogeneity would also favor the faithful reproducibility of culture and differentiation trajectories.

Forcing the formation of EBs using chemical and geometric molds can attenuate these diameter discrepancies. For example, the traditional hanging drop method allows small clusters of cells to aggregate through gravitational forces [13]. However, hanging drop methods also have their limitations, as EBs fabricated by the hanging drop method are generally small in size, have a low differentiation yield, and are prone to bursting/disaggregation [14]. The use of concave molds facilitates the aggregation of cells in a high-throughput and reproducible manner [15], and several reports have illustrated the use of ad hoc lab-made molds to produce spheroids (in general) and EBs (in particular).

Nevertheless, the use of commercially available culture devices has advantages over custom-made devices in terms of availability, reproducibility, and even price. For this reason, V-bottom well plates have been used extensively for the formation of EBs since their introduction. Now, coupled with an ultra-low-adhesion (ULA) layer, V-bottom plates have become the standard for EB formation [16]. Stem cells are prone to attach to surfaces, so an ultra-low adhesion coating is needed to allow the formation of cellular aggregates in well plates [17,18]. Therefore, the ULA V-bottom plates also enable a less labor-intensive EB fabrication and facilitate control over cellular aggregation [19,20]. However, these plates are still not readily available and can be expensive.

A number of methods have been reported to treat regular culture wells with different solutions and materials (i.e., agar or a polyvinyl alcohol (PVA) solutions) to confer them with anti-adherent properties [16,21]; however, these methods often involve multiple steps that can be cumbersome and time consuming. Here, we introduce and demonstrate the use of a cost-effective and rapid preparation method that imparts the ability to produce homogenous EBs at high yield and with a strong differentiation potential in conventional and commercially available non-ULA U-bottom and V-bottom well plates. Our method involves rinsing standard untreated tissue culture well plates with an anti-adherence solution 10 min before cell seeding, followed by centrifugation. This method is not resource-intensive and creates a hydrophobic surface that enhances cell aggregation in commercially available U- or V-bottom well plates for reliable formation of EBs that can efficiently differentiate into organoids.

## Methods

### Stem cell culture maintenance

The human embryonic stem cell (hESC) line H9 (WiCell, WA09) was obtained from the University of Leiden Medical Center. H9 cells were cultured at 37°C and 5% CO_2_ in feeder-free conditions in Essential 8™ (E8 medium (Gibco, Life Technologies, Carlsbad, CA, United States, cat. no. A1517001) in cell culture dishes coated with 5 μg/mL Vitronectin (Gibco, Carlsbad, CA, United States, cat. no. A31804) The hESCs were subcultured with 0.5 mM ethylenediaminetetraacetic acid (EDTA) (Invitrogen, Life Technologies, Waltham, MA, United States, cat. no. 15575020) every 4 days to avoid overcrowding and exponential differentiation. The passage number was kept below 60, and the cultures were routinely checked for mycoplasma contamination. Pluripotency was ensured by evaluating the expression of the markers Oct3/4 and SOX2.

### Well-plate coating

A 100 μL volume of anti-adherence rinsing solution (StemCell Technologies, Vancouver, Canada, cat. no. 07010) was added to untreated sterile CellStar U-bottom or V-bottom (Figure 1D) 96-well plates (Greiner Bio-One, Kremsmuenster, Austria, cat. no. 650185(U), 651161(V)). After incubation for 5 min at room temperature, the anti-adherence rinsing solution was aspirated, and the wells were washed with Dulbecco’s phosphate buffered saline (DPBS) (Gibco, Carlsbad, CA, United States, cat. no. 14190250) for another 5 min at room temperature. The DPBS was aspirated before cell seeding.

**Figure 1.**
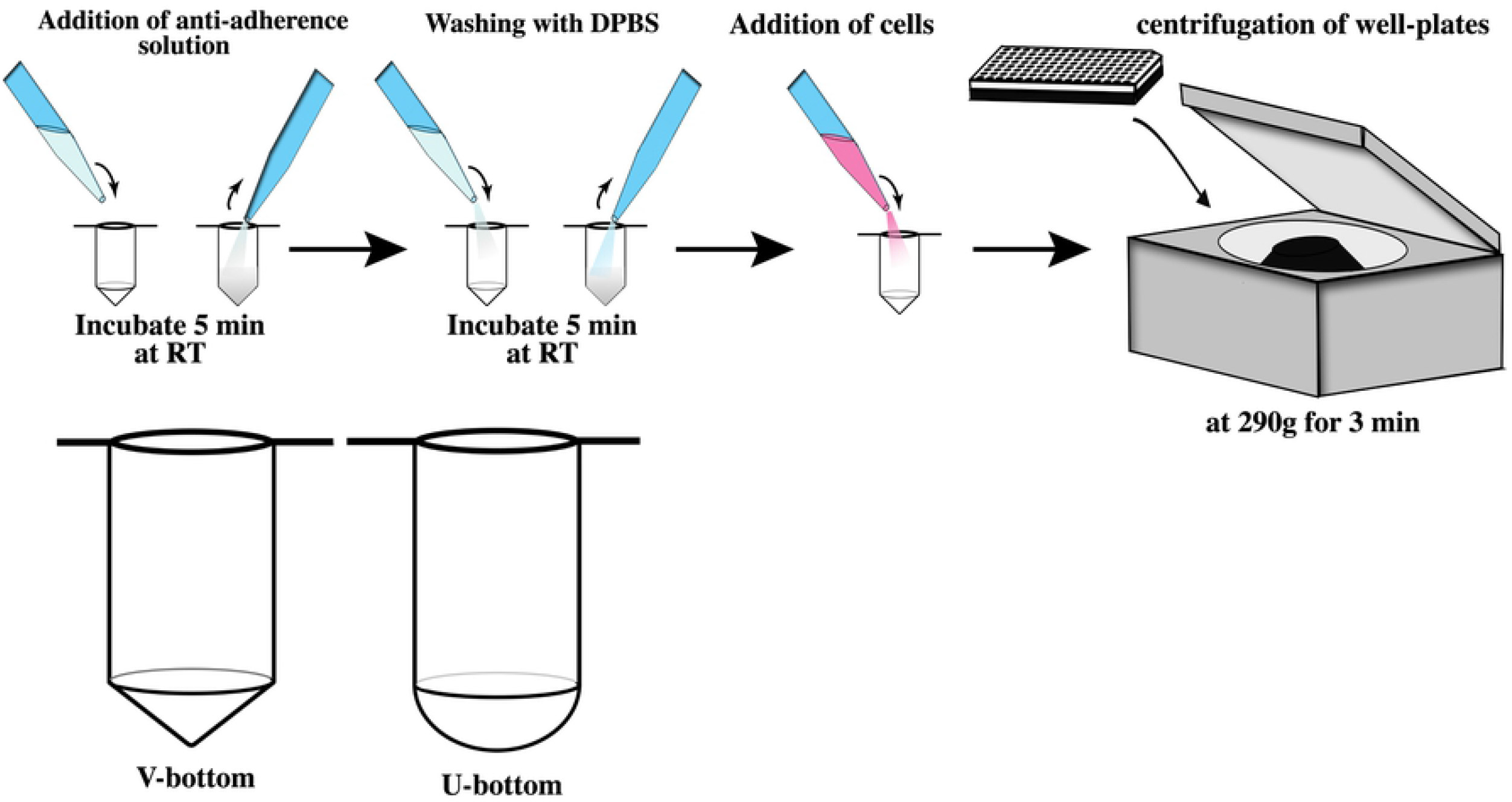
Protocol for embryoid body formation and different well geometries. We added anti-adherence solution into regular U- and V-bottom wells and incubated for 5 minutes at room temperature to develop anti-adherence coated plates and then washed gently with PBS. The application of an anti-adherence solution to conventional V-bottom and U-bottom plates enabled reproducible EB generation in low-cost well plates.

### Embryoid body formation

For EB formation, a previously published protocol was adapted [10] (Figure 1).

A single-cell suspension of hESCs was created by treatment of the cell culture with 0.5 mM EDTA for 3 min at room temperature. The released cells were collected, spun down, and counted. A range of 1,000–11,000 cells per well was seeded into wells prefilled with 150 μL Essential 6™ (E6 medium) (Gibco, Carlsbad, CA, United States, cat. no. A1516401) supplemented with 10 μM ROCK inhibitor (Selleckchem, Houston, TX, United States, cat. no. S1049). The well plates were then centrifuged at 290 × g for 3 min and incubated at 37°C with 5% CO_2_. After 24 h, the culture medium was changed to E6 medium supplemented with 2 μM XAV939 (Tocris Bioscience, Bristol, United Kingdom, cat. no. 3748), 10 μM SB431542 (R & D Systems, Minneapolis, MN, United States, cat. no. 1614/50), and 500 nM LDN193189 (StemCell Technologies, Vancouver, Canada, cat. no. 72147). The culture medium was changed daily.

### Microscopy and image analysis

Bright field images were collected every day using an EVOS cell imaging system (Thermo Fisher Scientific, Waltham, MA, United States). A 4× objective lens (16.0 mm working distance) was used for all images. Images were then imported into Adobe Photoshop CS6 (San Jose, CA, United States) for analysis. The EBs were isolated in the images by creating layer masks when contrasting an EB perimeter with its surroundings. The layer masks were smoothed using the “feather” function with a 5 pixel radius. Measurements (height, width area) and circularity were obtained using the “measurement” function of Photoshop.

### Organoid differentiation

Brain (telencephalic) organoids were differentiated using a previously published protocol [22] with minor modifications. In brief, 50 μL of Matrigel™ (3 mg/mL; Corning, New York, NY, United States, cat. no. 356230) was added to the wells of an untreated 96-well flat-bottom plate to create a bed layer and allowed to set for 15 min at 37°C. EBs were then transferred to individual bed layers and covered with an additional 50 μL of Matrigel to ensure full encapsulation of the EB. The top layer was allowed to set for another 15 min at 37°C, then differentiation medium was added. We conducted experiments of unguided and guided differentiation. In both cases, the basic differentiation medium consisted of Neurobasal-A (Gibco, Life Technologies, Carlsbad, CA, United States, cat. no. 21103049), 1× B27 without vitamin A (Gibco, Life technologies, Carlsbad, CA, United States, cat. no. 17504044), 1× GlutaMax (Gibco, Life Technologies, Carlsbad, CA, United States, cat. no. 35050038), and 1% (v/v) penicillin/streptomycin (Gibco, Life Technologies, Carlsbad, CA, United States, cat. no.15070063). For unguided differentiation, the medium was supplemented with 20ng/mL of FGF-2 (Gibco, Life technologies, Carlsbad, CA, United States, cat no. PHG0261) and 20ng/mL of EGF (Gibco, Life technologies, Carlsbad, CA, United States, cat no. PHG0311L). For guided differentiation, the basic differentiation medium was added with 3 μM CHIR99021 (Axon Medchem, Reston, VA, United States, cat. no. CT 99021) and 0.5 ng/mL bone morphogenic protein (BMP4, R & D Systems, Minneapolis, MN, United States, cat. no. 314). The medium was changed every other day. CHIR and BMP4 were only added for the first 3 days.

### Immunohistochemistry

After 30 days of culture, organoids were fixed in 4% formaldehyde for 60 min at room temperature, followed by an overnight incubation in 30% sucrose in PBS at 4°C. The organoids were then embedded in Shandon cryomatrix (ThermoFisher, cat. no. 6769006), flash frozen at - 8Ü°C, and cut into 5 μm sections using a cryostat (SLEE medical GmbH, Nieder-Olm, Germany; type MNT). After an additional fixation step with acetone at 4°C, the sections were blocked with 1% BSA and 0.025% Triton-X 100 in PBS for 1 h at RT.

The sections were then incubated with primary antibodies (rabbit anti-MAP2, 1:200, Abcam, cat.no. ab32454; mouse anti-GFAP, 1:500, Abcam, cat.no. ab212398; mouse anti-β3-tubulin, 1:500, Santa Cruz Biotechnology, cat.no. sc-80005) in blocking solution overnight at 4°C, followed by incubation with secondary antibodies and conjugated antibodies in blocking buffer (SOX2-GFP, 1:500, eBioscience, cat.no.53-9811-82; Donkey anti-mouse, AF647, ThermoFisher, cat. No. A31571, goat anti-rabbit AF594, Invitrogen, cat. No. R37117) for 1 h at RT. Nuclei were visualized with DAPI (ThermoFisher Scientific, cat.no. D1206) in PBS for 20 min.

### Image analysis and statistical analysis

The diameter of each EB was calculated using the average of the height and width of the spheroid. For comparison between variables, values were exported to Prism 8 (GraphPad, v 8.4.3, San Diego, CA, United States) for one-way analysis of variance (ANOVA) (P<0.0001) was used to establish significant differences (P<0.05).

## Results and discussion

Here, we demonstrated the use of standard, commercially available V-bottom and U-bottom well culture plates for the fabrication of functional neural EBs from human embryonic stem cells. Stem cells have a strong tendency to adhere to surfaces; therefore, we added a commercial anti-adherence solution to inhibit cell adhesion to the surface of wells and instead to favor cell aggregation. We also explored the use of a centrifugation step to further promote cell aggregation. Furthermore, we showed that the use of different initial cell seeding densities yielded EBs of different final diameters, thereby offering a way to precisely define and customize the size of neural EBS. This protocol yielded functional EBs that successfully differentiated into neural organoids.

Our integration of the use of a commercially available anti-adherence solution and a short centrifugation step resulted in a cost-effective and reproducible protocol for neural EB fabrication that can use conventional, readily available U- and V-bottom plates. The methods presently available for fabricating neural spheroids and EBs frequently depend on the use of ULA plates (both U- and V-bottom shaped) [23–25], which are more expensive and therefore less available than standard plates.

Our hESC culture system produced EBs after 6 days of culture. The EBs exhibited a characteristic neuroepithelial development, including a translucent body with a bright surface and smooth edges (Figure 2A,B) [10]. We used these (morphological) attributes for our initial assessments and to ensure the quality of EB formation. We found that EBs showing grainy, bumpy, or fading edges tended not to survive until the end of induction (day 6) (Supplementary S1 Fig.).

**Figure 2.**
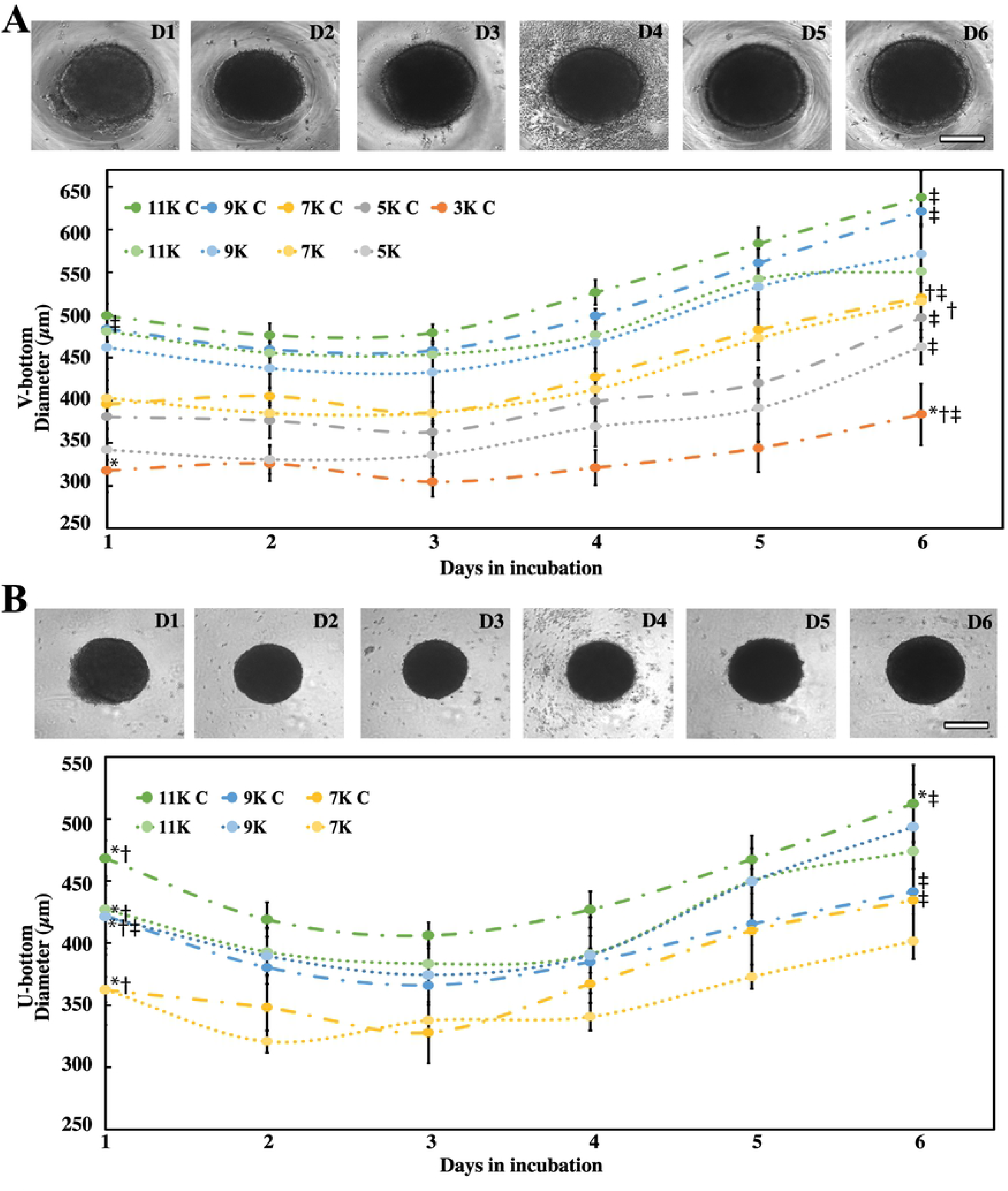
Characterization of embryoid body (EB) quality and daily growth. Examples of appropriate EB growth in V-bottom (A) and U-bottom (B) anti-adherence coated plates (AACPs) at different days (“D”). Graphs shows different color-coded seeding concentrations given in thousands (“K”) and if centrifuged (“C”). (A) The images depict EBs cultured in V-bottom AACPs for 6 days (D1–D6); their corresponding diameter is depicted in the graph below. Scale bar: 200 μm. EBs showed a characteristic neuroepithelial morphology, especially the white band along the inner border. (B) EB growth progression cultured in U-bottom AACPs. *P<0.001 indicates a significant difference for all EBs of the same well plate and centrifugation. †P<0.001 indicates a significant difference for EBs of the same well plate type but different centrifugation condition. ‡P<0.001 indicates a significant difference for EBs with different plate types but the same centrifugation condition. Scale bar: 200 μm. We considered at least 5 EBs (n=5) for each experimental group at the beginning of the experiment.

Cells in untreated well plates usually formed a central cluster, with small satellite aggregates along the slope of the well plate (S1 Fig.). We did not observe reproducible formation of consolidated tissue in wells not treated with the non-adherence solution. By contrast, most cells precipitated into a singular cluster in the treated wells (Figure 2A,B). Aggregation seemed to be positively correlated with survivability, as none of the EBs cultured in untreated well plates survived beyond the first couple of days. The inability to form single clusters resulted in loose cellular interactions and loss of the desired round EB shape. This presumably led to cell death and aggregate fragmentation (S1 Fig.).

### Effects of seeding density and geometry on EB size and shape

Figure 2A shows the size evolution of EBs cultured in V-bottom wells and derived from different initial cell counts per well. Trends associated with centrifuged and non-centrifuged plates are also compared. As expected, higher seeding densities (i.e., the number of cells seeded per well at the initial time point) resulted in higher initial and final diameters.

The EB diameter is influenced by the seeding concentration [26]. In our experiments, an increase in the diameter of the EBs during 6 days of culture was also a function of the initial cell density. In general, we observed an average increase of nearly 100 μm in the diameter of EBs formed at high initial cell densities (i.e., 9×10^3^ and 11×10^3^ cells per well). The EBs formed in the wells initially seeded with 5×10^3^ and 7×10^3^ cells per well showed an increase in diameter of nearly 75 μm, while the EBs formed at the lowest initial cell density of 3×10^3^ cells per well increased their diameter only modestly during the 6 days of culture. However, under equal seeding conditions, we did not observe significant differences between the average initial or final diameters of EBs when the cells were centrifuged or not centrifuged in the well systems (except for seeding densities of 9×10^3^ and 11×10^3^ cells per well).

Figure 2B shows the evolution of size in the EBs formed and cultured in U-bottom plates at different initial seeding conditions, with or without well centrifugation. Consistent with previous reports [27,28], our results suggest that the geometry of the well affects the process of EB formation. The V-bottom wells created EBs with a slightly larger (but not statistically significant) initial diameter than their U-bottom counterparts (Figure 2A). However, we observed clear differences in the final diameters of EBs formed in U-bottom versus V-bottom plates, as the EBs formed in V-bottom wells were significantly larger (Figure 2B and 3A). For example, EBs cultured for 6 days in U-bottom or V-bottom wells exhibited an average diameter of 456±13 and 525±44 μm, respectively (Figure 3A). Notably, the EBs generated in U-bottom plates only reached 500 μm in diameter at the highest seeding densities (i.e., 11×10^3^ cells per well) and with the assistance of the centrifugation step. The increase in diameter of the EBs from their initial size was also lower in U-bottom than in V-bottom plates. These observations can be explained using geometrical arguments, as V-bottom wells have concave bottoms with a steeper slope than U-bottom wells and therefore have a larger initial diameter. This means that more cells are able to precipitate to form larger agglomerates in V-bottom plates. Indeed, we observed that some cells remained floating in the culture medium in the rounder U-bottom wells and were not integrated to the main EB (Supplementary Figure 1F).

**Figure 3.**
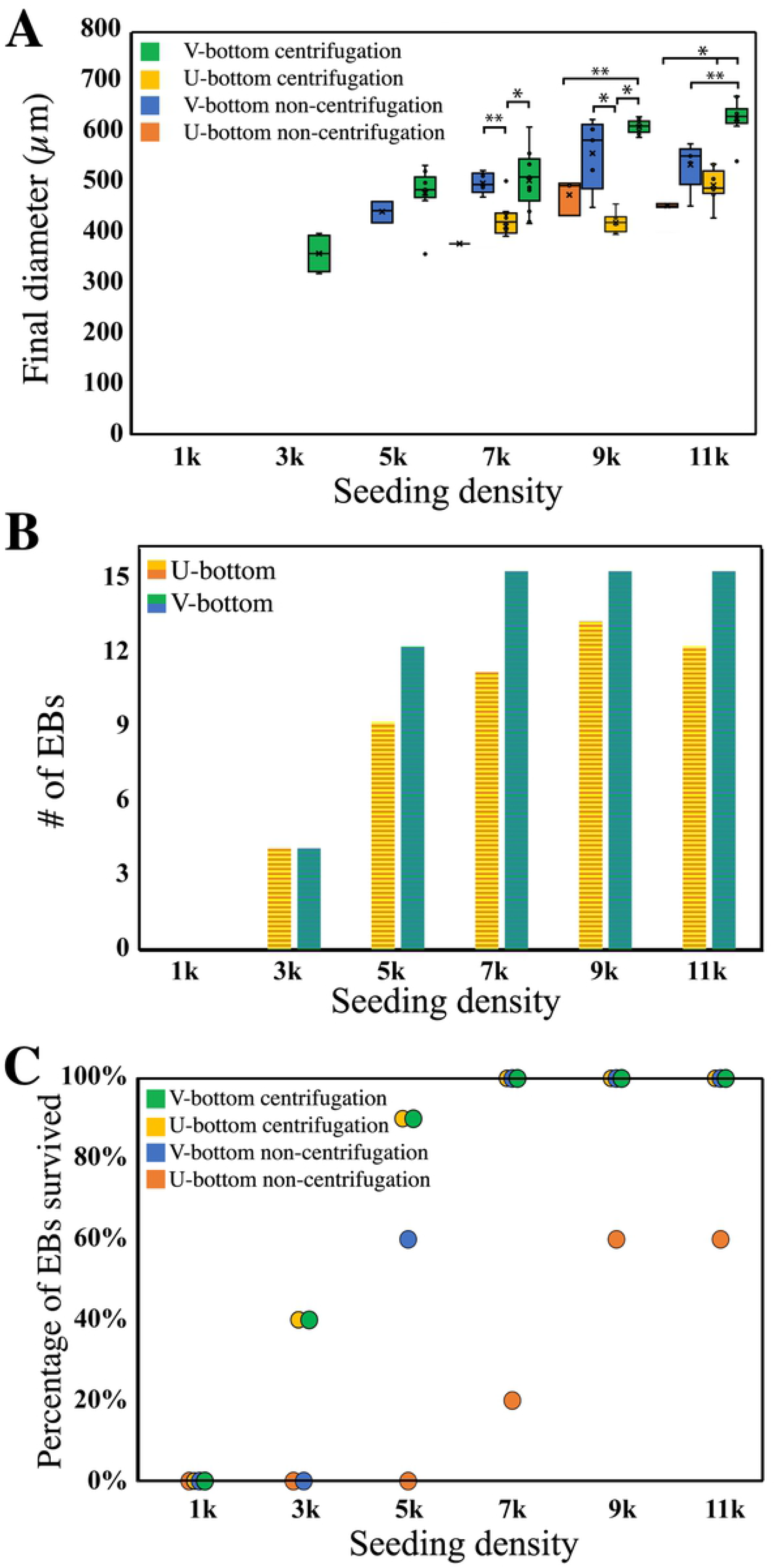
Diameter and survivability of EBs fabricated under different conditions. (A) Box-plot distribution of the final diameter of EBs at different concentrations. Boxes are color coded to indicate the type of well plate and centrifugation treatment used. Box plots include the maximum, minimum, and mean values observed. A single asterisk (*) indicates a significant difference of P<0.001. A double asterisk (**) indicates a significant difference of P<0.05. We evaluated at least 5 EBs (n=5) for each experimental group at the beginning of the experiment. Distribution of the diameters of EBs produced in V-bottom and U-bottom wells. Average diameters were determined based on the number of EBs (n) that survived the entire culture time. (B) Histogram of EBs produced in U-bottom and V-bottom wells that survived until the end of the experiment. EBs formed in U-bottom wells are shown in blue; EBs formed in V-bottom wells are represented in orange. The initial number of EBs formed = 15 in all experiments. (C) Percentage of the number of EBs that survived under the different fabrication conditions.

Figure 3A shows the average diameters achieved in all our EB-fabrication experiments. The largest EBs were obtained with V-bottom wells and centrifugation. Interestingly, the formation experiments in V-bottom wells indicated that the use of a centrifugation step has a clear effect only with high seeding densities (i.e., 9×10^3^ and 11×10^3^ cells per well). Figure 3A also shows the conditions that were most suitable for EB formation using U-bottom wells. In the absence of a centrifugation step, EB formation in U-bottom wells was feasible only at high seeding densities (i.e., equal or higher than 9×10^3^ cells per well). Centrifugation enabled the formation of EBs in U-bottom wells at a wider range of seeding conditions, from 7×10^3^ to 11×10^3^ cells per well. This was an improvement on previously reported methods that identified 9×10^3^ cells per well as a minimum threshold for reproducible EB formation [10,29,30].

Relevant indicators of the quality of the fabricated EBs were their final diameters and the span of their diameter dispersions. Previous research has shown that EBs with different diameters exhibit varying rates of differentiation or require different concentrations of induction factors to achieve homogeneous expression of differentiation genes [8,31]. Therefore, reducing the variability of EB diameters standardizes subsequent organoid differentiation. Figure 3A shows the experimental conditions that enabled the formation of more homogeneous sets of EBs.

Some of the experimental conditions tested resulted in homogeneous sets of EBs that exhibited diameters with standard deviations of less than 50 μm, which is an improvement on the tolerance level previously reported [9,10]. Indeed, the process of EB fabrication at high seeding densities (i.e., 9×10^3^ and 11×10^3^ cells per well) benefited greatly from the inclusion of a centrifugation step.

We also observed that EBs obtained using seeding densities of 9×10^3^ and 11×10^3^ cells per well did not differ significantly in their attributes (size and shape). This suggest the existence of an upper size threshold, of 620 μm in diameter, for the fabrication of EBs using V-bottom wells.

For example, EBs formed at high seeding densities in V-bottom plates exhibited a much lower dispersion in size with a centrifugation step than without. We also analyzed the sphericity (i.e., circularity and roundness) of the EBs formed under different fabrication conditions (Table 1). High values of EB circularity (i.e., above 0.8) resulted in size and shape homogeneity during subsequent culture and differentiation into organoids. We observed that our EBs had circularity exceeding 0.8 (Table 1), as reported previously for other EB protocols [32]. In general, the use of U- and V-bottom well plates did not affect EB circularity. The circularity was not significantly different at different cell seeding densities and remained constant throughout the culture.

**Table 1.** Circularity and roundness, as determined by image analysis of embryoid bodies (EBs) cultured in V-bottom and U-bottom wells at different seeding densities. All values were calculated after 6 days of induction.

| Seeding<br>(cells/well) | V-bottom |  | U-bottom |  | V-bottom |  | U-bottom |  |
| --- | --- | --- | --- | --- | --- | --- | --- | --- |
|  | no centrifugation |  | no centrifugation |  | centrifugation |  | centrifugation |  |
|  | Circularity | Roundness | Circularity | Roundness | Circularity | Roundness | Circularity | Roundness |
| 1,000 | - | - | - | - | - | - | - | - |
| 3,000 | - | - | - | - | 0.88±0.013 | 0.93±0.03 | - | - |
| 5,000 | 0.87±0.024 | 0.88±0.04 | - | - | 0.88±0.007 | 0.95±0.02 | - | - |
| 7,000 | 0.88±0.011 | 0.92±0.03 | 0.89±0.001 | 0.98±0.02 | 0.88±0.011 | 0.92±0.03 | 0.89±0.005 | 0.92±0.06 |
| 9,000 | 0.88±0.008 | 0.96±0.01 | 0.89±0.001 | 0.97±0.02 | 0.88±0.012 | 0.95±0.02 | 0.89±0.003 | 0.96±0.05 |
| 11,000 | 0.88±0.008 | 0.92±0.01 | 0.90±0.001 | 0.98±0.01 | 0.89±0.009 | 0.97±0.01 | 0.89±0.004 | 0.9±0.06 |

\*Values only provided if EB survived until Day 6.

### Effects of seeding density and geometry on EB survival

The survival of EBs differed depending on the fabrication conditions. Here, the term “survival” encompasses the maintenance of living cells and the retention of circularity, EB diameter, growth, and smooth edges (see S1 Fig. for exclusion criteria). Different cell seeding densities resulted in different percentages of EB survival. For both U-bottom and V-bottom wells, cell seeding densities of 1×10^3^ and 3×10^3^ cells per well did not yield viable EBs (Figure 1, Figure 3B,C, and Table 1). This suggests that a minimal initial number of cells (i.e., ~5×10^3^ cells) is needed (in the absence of centrifugation) to promote adequate cell aggregation and EB survival (Figure 3A, B). This is in line with previously reported protocols that also indicated a minimum culture seeding of 5×10^3^ cells for reproducible EB formation [33].

In general, we observed higher EB survival in V-bottom than in U-bottom wells (Figure 3B). For instance, in our experiments seeded at 5×10^3^ cells per well, we only observed EB survival when V-bottom wells were used. EBs seeded in both U- and V-bottom wells exhibited similar initial diameters at 5×10^3^ cell seeding. However, no EBs survived 6 days of culture in U-bottom wells, while EBs cultured in V-bottom plates had a survival rate of 60% (n=5) (Figure 3C). Similarly, a 100% survival was observed in experiments in V-bottom wells seeded at 7×10^3^, 9×10^3^, and 11×10^3^ cells per well, while the U-bottom wells yielded a 60% (n=5) survival (Figure 3C). The discrepancy in survival between U-bottom and V-bottom wells at similar seeding densities (and initial diameters) reinforces the importance of well-plate geometry in its ability to aggregate cells and form EBs (Figure 3B,C).

Previous experiments have shown the benefits of forced cell aggregation through centrifugation [14,34]. Centrifugation of well plates clearly aids in precipitating cells together, increasing their aggregation, cell-to-cell interactions, and survival. In our experiments, centrifugation immediately after cell seeding improved EB formation and survival at day 6, without affecting the initial EB diameter. EBs cultured in centrifuged V-bottom wells showed full (100%) survival at cell seeding densities of 7×10^3^ K, 9×10^3^ K, and 11×10^3^ K cells per well (Figure 3C). At seeding densities of 5×10^3^ K cells per well, survival was higher in centrifuged V-bottom wells than in non-centrifuged wells (90% versus 60%). Similarly, at seeding conditions of 5×10^3^ K cells per well, EBs survival was higher in centrifuged U-bottom wells (i.e., 90%) than in non-centrifuged U-bottom wells (i.e., 20%). Our results suggest that centrifugation is an important step in this protocol, as it increases the survival percentage of EBs at medium seeding densities.

### EB maturation into neural spheroids

We cultured EBs for extended time periods (i.e., more than 300 days) under guided [22] and unguided [35] differentiation protocols to induce them into telencephalic tissues and assess their functionality. All EBs formed in V-bottom and U-bottom wells showed budding tissue within the first couple of days of brain differentiation, which suggested their ability to develop into organoids once embedded into Matrigel (Figure 4A-D). Unguided differentiation involved supplementing the medium with 20ng/mL of EGF and FGF-2, growth factors known to induced maturation to neurons if the neural identity was secured during the induction process [36]. Guided differentiation involved the use of enriched media added with WNT (CHIR99021) and bone morphogenic protein (BMP4) activators, to expedite the dorsalization of the tissue and induce proper neural tube formation [22].

**Figure 4.**
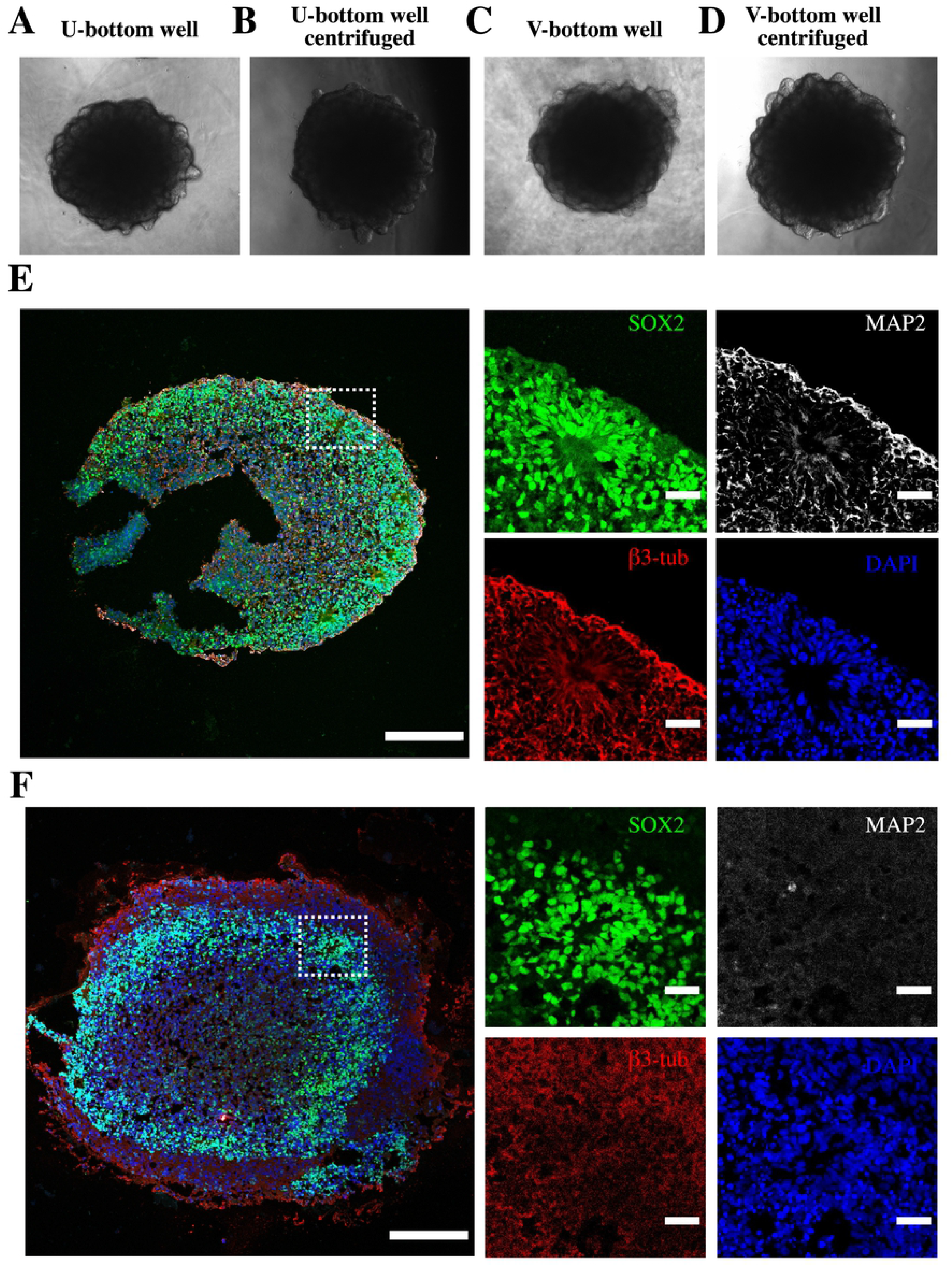
Examples of successful organoid differentiation from EBs. Brain organoids were formed using a guided and unguided differentiation protocols to induce telencephalic differentiation. Regardless of the use of U- or V-bottom AACP, the EBs demonstrated budding neuroepithelia at the periphery, hinting at successful organoid formation. Examples of EBs produced in a (A) a U-bottom well without centrifugation, (B) a U-bottom well with centrifugation, (C) a V-bottom well without centrifugation, and (D) a V-bottom well with centrifugation. Scale bar: 500 μm. (E-F) Successful organoid differentiation at 30 days of culture. Dotted lines indicate the rosette formations in the periphery of the organoid indicate the undifferentiated neuroepithelial marker SOX2 andthe neural identity markers of β3-tubulin and MAP2. EBs cultured under (E) guided (F) and unguided differentiation protocols showed similar structural location of the expressed markers, suggesting the neural predestination of the neuroepithelial tissue. Scale bar: 200 μm.

We observed proper EB induction into neuroepithelia in guided and unguided differentiation experiments; both types of experiments yielded rosette lumens, formations only seen in the neuroectoderm, a neural tube analog [37]. All EBs fully developed into rosette forming organoids after 30 days of culture and expressed the undifferentiated neuroepithelial marker SOX2 and mature neuronal markers of β3-Tubulin and MAP2 (Figure 4E,F). The abundant expression of SOX2 within these rosettes suggests the overall pluripotent nature of the tissue [38]. We observed lower expression of the MAP2+, an indicator of neuron maturity, in organoids produced by unguided than in those generated in guided differentiation protocols. Yet, the identical distribution of β3-Tubulin and MAP2 in the outer rim of the rosette suggested [24] a successful induction into neuroepithelia even in unguided differentiation experiments.

## Conclusions

The combination of using standard V-bottom plates pretreated with a commercial anti-adherent solution 5 min before cell seeding, coupled with a brief centrifugation step after seeding, results in a cost-effective and reliable fabrication method for producing homogeneous and functional EBs.

Although V-bottom wells generated larger EBs than U-bottom wells, we also identified experimental conditions that would enable the reliable use of U-bottom wells for EB fabrication. The centrifugation of the well-plate cultures in V- and U-bottom well plates effectively enhanced the survival and reduced the size variability of neural EBs. In general, the standard deviation in the size of EBs generated with centrifugation was smaller than 50μm, which is lower than the reported variability using other protocols (i.e., 100 μm) [9,10].

Although our method was standardized only for neuroepithelial EBs, we believe that it is translatable to the fabrication of a wide spectrum of organoids representative of different tissues. Many organoid cultures begin with circular EBs, including those related to kidney [26], liver [39], heart [40], optic cup [1], and other organs and tissues [38,41]. The methods provided here could potentially facilitate and streamline organoid cultures in a diverse spectrum of labs spanning different fields.

The protocols introduced here are simple and easily translatable to any cell culture laboratory. Their simplicity and cost-effectiveness may facilitate the entry of more laboratories into the nascent field of organoid fabrication and organoid-based research.

## Acknowledgments

We thank Carla Cofiño Fabres for providing V-bottom well plates. We also thank Mariana Garcia-Corral and Carla Annink for their support in cell culture techniques and standardization.

## Supporting Information

**S1 Fig. Examples of poorly formed EBs at different stages of induction**. (A) Low seeding concentrations or non-coated wells produce small cell aggregates. These small aggregates will not fuse together later and will not develop into EBs. Scale bar: 200 μm. (B) Central cluster with satellite aggregates in non-coated wells. As in (A), these peripheral aggregates will not fuse with the main cluster. Scale bar: 200 μm. (C) Disintegrated EB; remains of an EB that did not survive until the end of the experiment. Scale bar: 200 μm.

## Notes

### Competing Interest Statement

The authors have declared no competing interest.

